# Transthyretin orchestrates vitamin B12-induced stress resilience

**DOI:** 10.1101/2024.02.07.578164

**Authors:** G. Stein, J.S. Aly, A. Manzolillo, L. Lange, K. Riege, I. Hussain, E.A. Heller, S. Cubillos, T. Ernst, C.A. Hübner, G. Turecki, S. Hoffmann, O. Engmann

## Abstract

Chronic stress significantly contributes to mood- and anxiety disorders. Previous and current data suggest a correlative connection between vitamin B12 supplementation, depression, and stress resilience. However, the underlying mechanisms are still poorly understood. This study reveals a molecular pathway that functionally connects vitamin B12, DNA methylation (DNAme), and stress resilience. We identified Transthyretin (*Ttr*) as a sex-specific key target of vitamin B12 action in chronic stress. Accordingly, *TTR* expression was increased postmortem in the prefrontal cortex of male, but not female, depressed patients. Virally altered *Ttr* in the prefrontal cortex functionally contributed to stress- and depression-related behaviors, changes in dendritic spine morphology and gene expression. In stressed mice, vitamin B12 reduced DNAme in the *Ttr* promoter region. Importantly, using *in vivo* epigenome editing to alter DNAme in the brains of living mice for the first time, we establish a direct causal link between DNAme on *Ttr* and stress-associated behaviors. In summary, using state-of-the-art techniques, this study uncovers a mechanistic link between cobalamin supplementation and markers of chronic stress and depression, encouraging further studies into environmental interventions for mood disorders.

## Introduction

Chronic stress is a major risk factor for mental illnesses, such as major depressive disorder (depression, MDD), which ranks among the leading causes of disability and suicide ^1,2^. However, the mechanisms by which chronic stress impacts mental health, and how consequences can be ameliorated by rapid environmental interventions such as dietary factors, remain poorly understood ^3^.

Vitamin B12 (cobalamin) is an essential nutrient synthesized by bacteria. It serves as a cofactor in the one-carbon metabolism, which provides methyl donors for the methylation of DNA and proteins, including histones. Notably, a common symptom of vitamin B12 deficiency is a depressive phenotype ^4^ and individuals deficient in vitamin B12 have an elevated risk of developing depression ^5–9^. Moreover, there is evidence that vitamin B12 supplementation in non-deficient populations may lower the risk of depression ^10–12^, and help to reverse the pro-depressive effects of early life stress in animal models ^13^. Previous research in our team demonstrated that a single acute dose of vitamin B12 reduced depressive-like behaviors in mice in the chronic mild stress (CMS)-model ^14^. Additionally, vitamin B12 rescued stress-induced alterations in plasma mannose and Gmppb ^15^, a key enzyme of glycosylation metabolism associated with depression ^16^. Despite these correlative findings, it is unclear how vitamin B12 mediates stress resilience and improves markers of depression on a molecular level.

Here we explored the effects of an acute dose of vitamin B12 on stress response using a multilevel approach. To that end the chronic variable stress (CVS) mouse model allowed a sex-specific analysis that closely mirrors the molecular aspects of human depression ^17^. Through next-generation RNA sequencing, we identified Transthyretin (*Ttr*) as a pivotal stress-regulated gene product, which was specifically rescued by vitamin B12 in the prefrontal cortex (PFC) of male mice. Accordingly, we observed altered *TTR* levels in the postmortem PFC of male depressed patients. *Ttr*, a clinically relevant carrier protein for thyroid hormones and retinol, has been previously found to be altered in rodent models of stress ^18,19^ and is increasingly recognized for its contribution to various brain functions ^20^. However, modes of reversing *Ttr* levels remain elusive. *TTR* missense mutations are linked to hereditary amyloidosis, a rare and lethal disease. In this context, *TTR* stands as the first gene-edited by CRISPR technology *in vivo* in humans ^21^. Yet, despite its pivotal clinical role, surprisingly little is known about the chromatin regulation of the *Ttr* gene.

By using a viral approach to alter *Ttr* levels in the PFC, we establish a causal connection between *Ttr*, dendritic spine morphology, and stress-associated behaviors, and we determine transcriptional pathways downstream of *Ttr*. Furthermore, by pioneering the combined use of DNAme editing and jetPEI *in vivo* transfection strategies, we were able to alter the *Ttr* chromatin state in the brains of living mice. We observed that DNAme editing on distinct loci in the *Ttr* promoter region caused behavioral changes associated with stress and mood. In summary, these data unveil a sex-specific molecular pathway with the potential to rapidly ameliorate depression-associated phenotypes and increase stress resilience.

## Methods

Extended methods are found in the supplementary material.

### Animal housing and experimental approval

Mice were accommodated in accordance with ethical guidelines set by the Thüringer Landesamt für Verbraucherschutz (TLV). All experiments were carried out under the approved animal license UKJ-18-037 (Germany), aligning with the guidelines of EU Directive 2010/63/EU for animal experimentation. The studies involving genetically modified organisms adhered to S1 regulations as outlined in the GenTAufzV. C57Bl/6J mice were bred at the animal facilities of the University Hospital Jena, Germany, and Friedrich-Schiller-University, Jena, Germany. The mice were provided with standard chow (LASQCdiet Rod16-R, LASvendi GMBH, Soest, Germany), containing 50 mg/kg of vitamin B12. The mice used in the experiments were a minimum of 10 weeks old, with both sexes utilized as specified. Housing conditions involved a 14L:10D light cycle.

### CRISPR-dCas9 epigenome editing

Vectors coding the dCas9-DNMT3ACD-DNMT3LCD-3xFLAG fusion gene (contains the enzymatic group of the DNA methyltransferase DNMT3) and the control plasmid pCMV-dCas9-mD3A (#78257, Addgene, Watertown, MA, USA) were used as described in ^14,22^. Guide-RNAs against the promoter region of *Ttr* were designed via a published protocol (available at www.addgene.org/crispr/church/). Scrambled controls were used as described ^14,22^.

### Cell culture and transfection

Neuro2A-cells (ATCC #CCL-131) were cultured in six-well plates (10^6^ per well) in DMEM + GlutaMAX-1 (25300-054/096, GIBCO by Thermo Fisher Scientific, Waltham, MA, USA) with 10 % fetal bovine serum (South American Origin, #51810-500, Biowest) and 1 % Penicillin-Streptomycin (#15070-063, GIBCO). For CRISPR epigenome editing *in vitro*, vectors were co-transfected using Lipofectamine 2000 transfection reagent (#11668019, ThermoFisher Scientific) in OptiMEM (#31985070, GIBCO). The medium was replaced after 4 h with DMEM + GlutaMAX-1 medium. 48 h after transfection, cells were collected for DNA or RNA extraction.

### JetPEI *in vivo* transfection

The procedure was essentially performed as described ^23^. 12 µg of plasmid DNA (ratio 1:10 for guide-RNA and dCas9-construct containing plasmids) was diluted into 5 µl of sterile 10 % glucose and added to diluted *In vivo*-jetPEI® reagent (#101000040 VWR, Polyplus-transfection). The solution was mixed by pipetting and incubated at RT for 15 min. A total of 1.5 µl of jetPEI/Plasmid mix was delivered into the PFC at a rate of 0.2 µl per min, followed by 5 min of rest. Mice were tested 2 days later.

### Drugs and chemicals

Mice were intraperitoneally (i.p.) injected with 2.7 mg/kg vitamin B12 (cyanocobalamin, #V6629, Sigma) or saline at an injection volume of 10ml/kg body weight and tested 24 h later.

### RNA purification and quantification

RNA extraction involved resuspension in Trizol and chloroform precipitation, followed by washing with isopropanol and 75% ethanol. Subsequently, cDNA conversion was performed using the GoScript™ Reverse Transcriptase kit (#A5001, Promega). Quantitative real-time PCR was conducted on a Bio-rad CFX96 Real-time system. The primer sequences are provided in the supplementary material. The quantitative PCR (qPCR) results were analyzed as described ^14^.

### AAVs & stereotaxic surgery

Bilateral stereotaxic surgery targeting the PFC was conducted following established procedures ^17^. The following three viruses were employed: pAAV.1-CAG-GFP (#37825, Addgene), OE-Ttr-GFP: pAAV-CAG-GFP-P2A-Ttr-WPRE.bgH, KD-Ttr-GFP: pAAV-U6-Ttr-shRNA-2-CAG-GFP-WPRE3 (Ttr-AAVs custom-made by Charitè viral vector core, Berlin, Germany).

### Behavioral tests and CVS

The acute behavioral tests tail suspension test, forced swim test, sucrose preference, splash test, and novelty-suppressed feeding were conducted as outlined in previous studies ^17,24^. The sucrose ratio was calculated as the amount of consumed sucrose divided by the amount of total consumed liquids. To assess anxiety, the open field test was used as described ^14^. No tail suspension test was conducted after CVS, as the tail suspension is part of the stress-induction protocol. Within CVS groups, vitamin B12 injections were administered on the last day (day 21), just before the stressor. The CVS protocol comprised 21 days of stress with alternating stressors (tube restraint, tail suspension, or 100 mild electric random foot shocks for 1 h each) ^17^. To mitigate variability related to the sex-specific scent of experimenters, a used male t-shirt was wrapped in clean protective clothing and placed in the experimental room during experiments ^25^. All experiments were conducted during the light phase of the light cycle to ensure comparability with prior experiments ^14^.

### Dendritic spine analysis

The analysis relied on the detection of AAVs’ GFP-fluorophores. Images of 40 µm PFA-fixed brain sections were captured using a Zeiss LSM 880 confocal microscope employing the AiryScan method. Maximum intensity projections were generated using Zen Black and Zen Blue software and subsequently analyzed in NeuronStudio (CNIC, Mount Sinai School of Medicine). The overall density of spines, proportions of thin, mushroom, and stubby spines, along with cumulative neck length, were determined using Graph Prism. The density of neck-containing spines refers to the sum of thin and mushroom spine densities.

### Next-generation RNA-sequencing

Eukaryotic mRNA sequencing was conducted by Novogene (U.K.) and by the sequencing core facility of the Fritz-Lipmann-Institute for Aging, Jena, Germany. An in-house RNA-sequencing analysis pipeline, described at https://github.com/Hoffmann-Lab/rippchen, was applied. Read quality trimming was performed using Trimmomatic ^26^ v0.39 (5nt sliding window, mean quality cutoff 20). Clipping off the Illumina universal adapter and removal of poly mono-nucleotide content from the 3’ reads end were done according to FastQC v0.11.9 reports using Cutadapt ^27^ v2.10. Sequencing errors were recognized and corrected with Rcorrector ^28^ v1.0.4 and ribosomal RNA-derived sequences were filtered using SortMeRNA ^29^ v2.1. The preprocessed data was aligned to the reference genome GRCm38 (mm10) using Segemehl ^30,31^ v0.3.4 in splice-aware mode with an accuracy cutoff of 95%. Unambiguously aligned reads were deduplicated for over-amplified PCR fragments based on unique molecular identifiers utilizing UMI-tools ^32^ v1.1.1 and quantified on Ensembl v102 reference annotation via featureCounts ^33^ v2.0.1 (exon-based meta-feature, minimum overlap 10nt). Experiment strandedness was inferred using RSeQC ^34^ v4.0.0. Significantly differentially expressed genes and their enrichment for biological pathways were determined using DESeq2 ^35^ v1.34.0, clusterProfiler ^36^ v4.2.0 with fGSEA ^37^ respectively. Changes with an adjusted P-value < 0.05 were considered for further analysis.

### DNAme analysis

DNA was purified and bisulfite converted as described in ^14^ using the DNeasy Blood & Tissue kit (#69506, Qiagen) and the EZ DNA Methylation-Gold kit (#D5006, Zymo, Irvine, CA, USA). PCR was performed with a Gotaq G2 Hot Start Polymerase system (#M7405, Promega). Pyrosequencing was conducted on a Pyromark Q96 Sequencer using matching reagents (#97204, #978703 Qiagen).

### Postmortem samples

Samples were generously provided by the Douglas-Bell Canada Brain Bank. Experiments were conducted in agreement with Douglas-Bell Canada Brain Bank (REB Approval #04/21) and the Ethics Committee of Jena University Hospital, Germany (Reg.-Nr. 2020-1862-Material). Groups were balanced for age and postmortem interval. Samples with postmortem interval > 100 h were excluded.

### Statistics

Statistical analyses were performed utilizing GraphPrism. The comparison of two groups with equally distributed variances employed a two-tailed Student’s t-test. In case of unequal variances, Welch’s t-test was used. One-way ANOVA and Tukey *post hoc* test were applied when varying a single factor across multiple groups, while Two-way ANOVA and Bonferroni *post hoc* test were employed when two factors were varied. The cumulative head diameter of dendritic spines was analyzed using the Gehan-Breslow-Wilcoxon test ^38^. Data points more than two standard deviations away from the average were considered outliers and subsequently removed. During behavioral tests and dendritic spine analysis, experimenters were blind to group assignments to minimize bias.

## Results

### Vitamin B12 reverses stress-induced changes in *Ttr*-expression

We hypothesized that the vitamin B12-induced reversal of stress effects may be, at least in part, mediated by transcriptional alterations given its’ putative influence on DNAme and histone methylation. To investigate this, we conducted RNA sequencing on the PFC using the CVS mouse model (**Fig. 1A**, **B**).

**Fig. 1:**
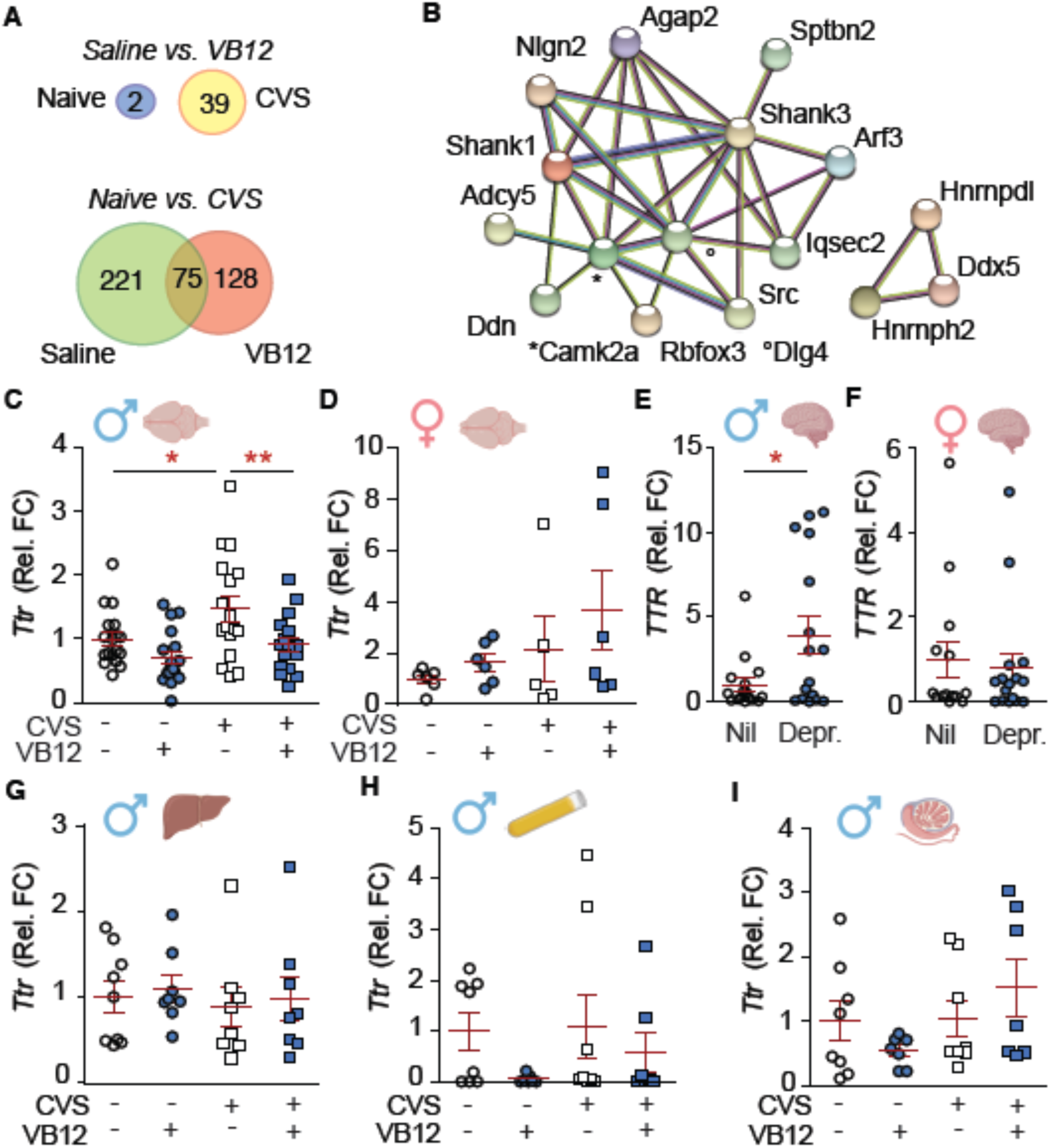
CVS and depression affect transthyretin in a sex- and tissue-specific manner and vitamin B12 reverses stress-effects in mice. **A**) Venn diagram of RNA-sequencing (males). **B**) Gene expression clusters that were differentially expressed by VB12 within the CVS group (string-db network). **C**-**F**) qPCR on PFC. **C**, **D**) Validation of RNA-sequencing data in an independent cohort of mice confirms reversal of stress-induced *Ttr*-changes by vitamin B12 (VB12) in males. **C**) Males: n = 17-18; stress effect: F(1,66) = 6.88, P < 0.05; drug effect: F(1,66) = 10.62, P < 0.01; *post hoc* test: stress within sal: *P < 0.05; drug within VB12: **P < 0.01. **D**) Females: n = 5-6; stress effect: F(1,19) = 2.62, P = 0.12; drug effect: F(1,19) = 1.25, P = 0.28. **E**, **F**) *TTR* is increased in postmortem samples of male but not female depressed patients (Depr.) vs. controls (Nil). **E**) Males: n = 15-16; Welch’s t-test: t_18.74_ = 2.47, *P < 0.05. **F**) Females: n = 14-16; t_28_ = 0.36, P = 0.72. **G**-**I**) qPCR on other mouse tissues reveals no effect of CVS or VB12. **G**) Liver: n = 8-9; stress effect: F(1,29) = 0.32, P = 0.58; drug effect: F(1,29) = 0.23, P = 0.64; interaction: F(1,29) < 0.001, P = 0.98. **H**) Plasma: n = 7-8; stress effect: F(1,26) = 0.53, P = 0.47; drug effect: F(1,26) = 2.81, P = 0.11; interaction: F(1,26) = 0.26, P = 0.61. **I**) Testes: n = 7-8; stress effect: F(1,26) = 2.73, P = 0.11; drug effect: F(1,26) = 0.02, P = 0.97; interaction: F(1,26) = 2.38, P = 0.14. **C**-**I**) Individual data points are plotted and means **±** s.e.m. are shown. Non-significant comparisons not unless specified. Sketches were generated with biorender.com.

Pathways that were abundant in differentially expressed genes within the CVS group included chromatin regulation, synaptic and intracellular signaling, metabolism, RNA regulation, and protein processing, indicating a widespread restructuring of the PFC in response to stress (**Table S1**). Well-established depression-associated gene products, such as *Fkbp5*, *Fosb*, and *Per1*, were among the affected genes in the CVS group. Additionally, CVS altered the expression of the folate receptor *Folr1*, the methyl transferases *Kmt5a*, *Tmt1A*, *Hnmt*, *Lcmt2,* the methyl transferase reductase *Mttr*, and the demethylase *Alkbh5*, suggesting more profound stress effects on methyl group metabolism (**Table S1**).

Vitamin B12 induced minimal transcriptional changes in the PFC of naïve mice. However, it partially reversed the transcriptional changes induced by CVS (**Fig. 1A, Table S1**). Among those genes differentially expressed by vitamin B12 within the CVS group were several key synaptic proteins including *Camk2a* and the autism-associated genes *Shank1*, *Shank3*, and *Nlgn2* (**Fig. 1B**).

*Ttr* expression was increased by CVS and rescued by vitamin B12 within the CVS group. Other genes exhibiting a similar pattern included *p35* (*Cdk5r1*, a Cdk5-activator implicated in learning, memory, schizophrenia, and neurodegeneration ^42,43^), the RNA binding proteins *Hnrnpdl* and *Hnrnph2*, the tyrosine kinase *Src*, the cytoskeletal regulator *Trak1* and the E3 ubiquitin-protein ligase *Znrf1* (**Table S1**).

The recurrent association of *Ttr* levels with stress in mouse PFC indicates biological robustness in the association between *Ttr* and chronic stress ^18,19^. However, as there is no known mechanism of reversal, we chose to focus our subsequent investigations on *Ttr* in order to characterize modes of regulating this pivotal gene and, in consequence, to reverse states that contribute to pathology.

The CVS-induced increase in *Ttr* and its reversal by vitamin B12 were confirmed by qPCR in an independent cohort (**Fig. 1C**). In female mice, neither CVS nor vitamin B12 exerted an effect on *Ttr*-levels in the PFC (**Fig. 1D**).

Next, we examined whether *TTR* levels were affected postmortem in the PFC of depressed patients and suicide victims. In male depressed patients, but not in females, *TTR* levels were elevated (**Fig. 1E, F**). *TTR* was particularly increased in male depressed patients who died from accidents or natural causes and less increased in suicide victims (**Fig. S1A**, **B**). In females, neither cause of death nor mental health status significantly affected *TTR* levels (**Fig. S1C**).

TTR functions as a carrier protein freely circulating in the cerebrospinal fluid of the brain and the bloodstream ^20^. Therefore, we assessed *Ttr* levels in other tissues. The liver has a strong baseline expression of *Ttr*. However, neither CVS nor vitamin B12 affected *Ttr* levels in the liver of male mice (**Fig. 1G**). In the brain, *Ttr* is secreted into the cerebrospinal fluid, which enters the systemic blood circulation by intracranial venous sinuses or cervical lymphatics. Moreover, *Ttr* can pass the blood-brain barrier ^44^. Nevertheless, neither CVS nor vitamin B12 had an impact on *Ttr* levels in plasma (**Fig. 1H**). Finally, we assessed testes tissue. Chronic stress can affect certain health measures in the offspring through the patriline (male germline) ^45^. The molecular mechanisms that transfer stress, a subjective effect originating in the brain, towards the germline are still unclear. Since *Ttr* can pass the blood-brain barrier, it is conceivable that it may enter the testes as well. However, *Ttr* levels were not increased in the testes in our data set (**Fig. 1I**). In summary, these data suggest that the observed CVS-induced increase in *Ttr* is specific to the brain, limited to the male sex, and can be rapidly reversed by vitamin B12.

### Viral-mediated alterations in PFC-*Ttr* alter depressive-like behaviors and dendritic spine morphology

Next, we investigated whether *Ttr* plays a causal role in the behavioral and morphological changes associated with chronic stress, given that previous associations between *Ttr* and stress had been largely correlative ^18,19^. To address this, we stereotaxically injected AAVs to overexpress (OE) or knockdown (KD) *Ttr* within the PFC of male mice. The AAVs independently expressed GFP, facilitating dendritic spine analysis. Analysis was conducted not earlier than 4 weeks post-injection to ensure stable expression of the constructs. The AAVs had no discernable impact on body weight or behavior in the open field (**Fig. S2**).

OE-*Ttr* elicited a robust increase in depressive-like behaviors across a variety of tests linked to escape behavior (tail suspension test), grooming (splash test), and feeding in a novel environment (novelty suppressed feeding) (**Fig. 2A-E**). Additionally, OE-*Ttr* led to a reduction in the density of stubby and neck-containing spines, mirroring the observed effects of CVS (**Fig. 2H-K, Fig. S3**). However, in contrast to CVS, OE-*Ttr* resulted in a decrease in the cumulative head diameter.

**Fig. 2:**
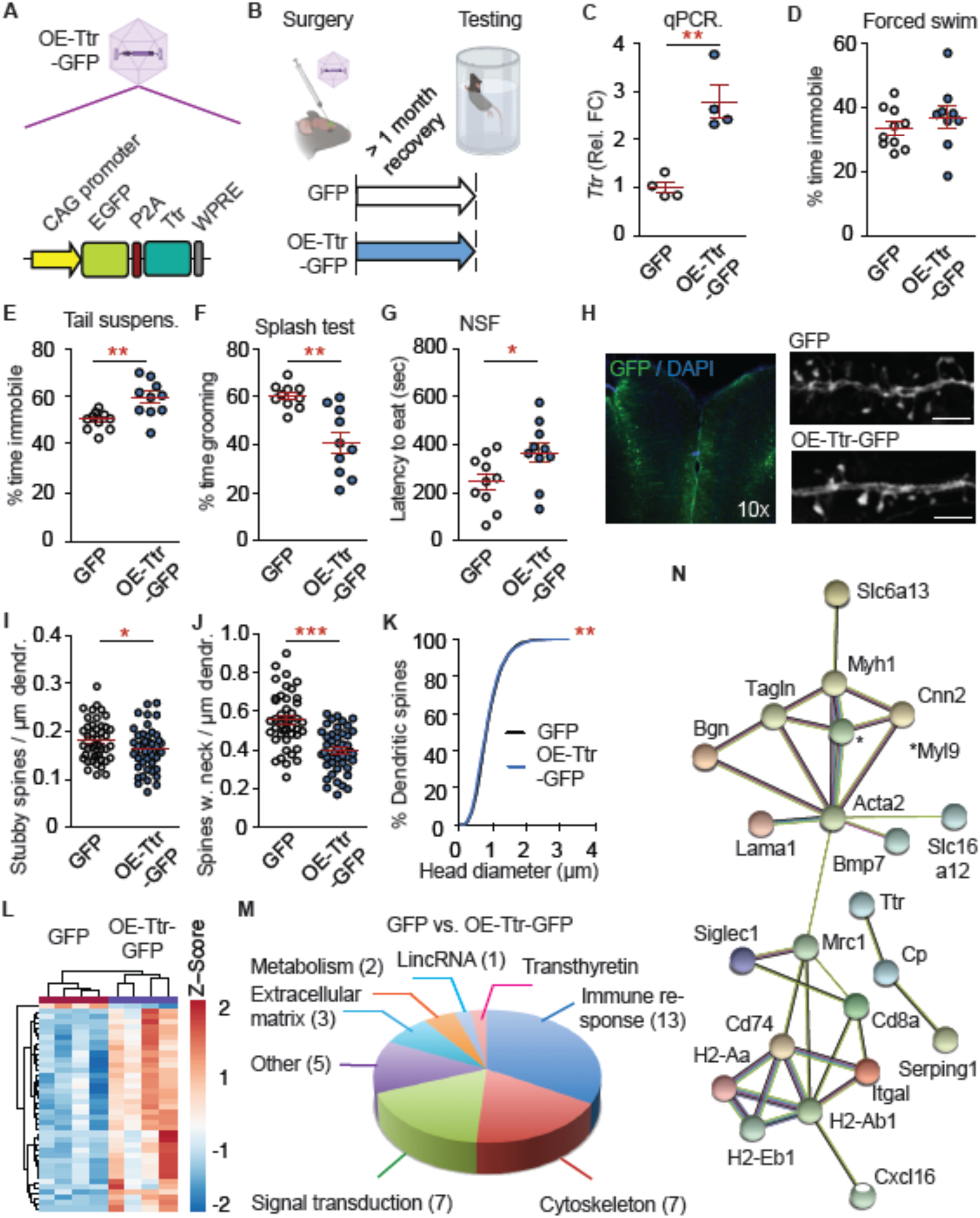
Overexpression of *Ttr* mimics effects of chronic stress and alters gene products for synaptic outgrowth. **A**) Schematic of the OE-Ttr-GFP AAV. **B**) Experimental plan. **C**) *Ttr* levels are increased by the OE-AAV. n = 4; t_6_ = 5.02, **P < 0.01. **D**-**H**) Overexpression (OE) of *Ttr* evokes depressive-like behaviors. **D**) Forced swim test: n = 9-10; t_17_ = 0.93, P = 0.37. **E**) Tail suspension test: n = 10; t_18_ = 3.44, **P < 0.01. **F**) Splash test: n = 10; Welch’s t-test: t_11.31_ = 4.14, **P < 0.01. **G**) Novelty-suppressed feeding (NSF): n = 10; t_18_ = 2.26, *P < 0.05. **H**-**K**) OE-*Ttr* mimics chronic stress effects on dendritic spines. **I**) Overview of OE-AAV infection in the PFC and representative images of dendrites. Scale bar 10 µm. **I**) Stubby spines: n = 43-46 dendrites from 5-6 mice; t_87_ = 2.01; *P < 0.05. **J**) Neck-containing spines: n =43-47 dendrites from 5-6 mice; t_88_ = 5.94; ***P < 0.0001. **K**) Cumulative head diameter: χ^2^ = 7.37, df = 1, **P < 0.01. **L**-**O**) RNA-sequencing after viral OE of *Ttr*. **L**) Heatmap of differentially expressed genes. **M**) Pie chart of assigned pathways. **N**) String-db analysis of shared pathways (RNAs with at least 1 connection shown). The cluster in the upper part is linked to cytoskeleton and synaptic growth. **C**-**G**, **I**, **K**) Individual data points are plotted and means **±** s.e.m. are shown. Sketches were generated with biorender.com.

RNA-sequencing revealed that OE-*Ttr* altered transcripts with a variety of functions, including immune response (*Cd74*, *H2-Aa*, *H2-Ab1*, *H2-Eb1*, *Cd8a*, *Ifitm1*, *Igha*, *Ighg2b*, *Igkc*, *Tgtp1*), carbohydrate metabolism (*Mrc1*, *Igfbp2*), and signal transduction (*Serping1*, *Slc6a13*, *Slc6a20a*, *Slc6a20a*) (**Fig. 2L-N, Table S2**). Several gene products were linked with either the cytoskeleton or the extracellular matrix (*Tagln*, *Bgn*, *Cnn2*, *Myh11*, *Acta2*, *Myl9*, *Lama1*, *Cilp*), suggesting an interplay of intra- and extracellular mechanisms that could influence dendritic spine outgrowth and synaptic plasticity. Taken together, OE-*Ttr* mimics specific behavioral and morphological aspects of chronic stress, suggesting that *Ttr* functionally contributes to stress symptoms.

Next, we asked whether a reduction of *Ttr* could counteract specific CVS effects as was observed with vitamin B12 (**Fig. S3**). Consequently, KD-*Ttr* effects were assessed under stress conditions. KD-*Ttr* blocked CVS effects on escape behavior (forced swim test; tail suspension was not applicable due to the CVS protocol), grooming (splash test), and novelty-suppressed feeding, but not sucrose preference (**Fig. 3A-G, Fig. S6**). This behavioral profile was consistent with OE-*Ttr*. In line with the preceding data, KD-*Ttr* rescued CVS effects on stubby and neck-containing spines and may have partially prevented CVS effects on the cumulative head diameter (**Fig. 3 H-N**). In summary, these findings suggest that increased *Ttr* contributes to behavioral and spine-morphological changes associated with chronic stress, while decreased *Ttr* can, at least in part, mitigate the effects of stress.

**Fig. 3:**
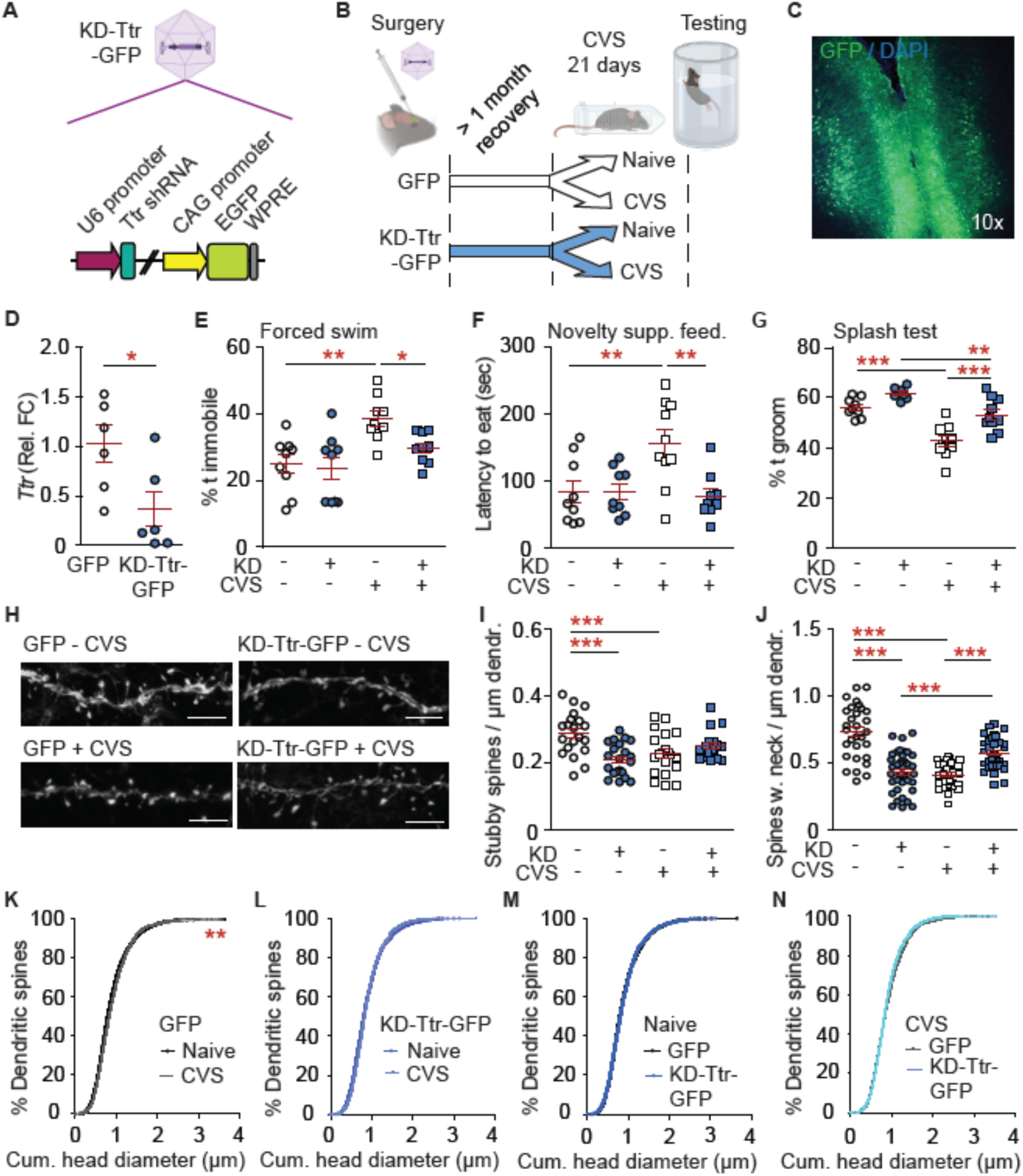
Knockdown of *Ttr* blocks effects of chronic stress. **A**) Schematic of the KD-Ttr-GFP AAV. **B**) Experimental plan. **C**) Overview of KD-AAV infection in the PFC. **D**) qPCR: n = 6; t_10_ = 2.52, *P < 0.05. **E**-**H**) Behavioral tests show a reversal of stress-induced behavioral changes by KD of *Ttr*. **E**) Forced swim test: n = 9; stress effect: F(1,32) = 14.60, P < 0.001; *post hoc* test: stress effect within GFP: **P < 0.01; AAV within CVS: *P < 0.05. **F**) Novelty suppressed feeding: n = 9-10; stress effect: F(1,33) = 4.38, P < 0.05; AAV effect: F(1,33) = 6.36, P < 0.05; interaction: F(1,33) = 6.43, P < 0.05; *post hoc* test: stress within GFP: **P < 0.01; AAV within CVS: **P < 0.01. **G**) Splash test: n = 8-10; stress effect: F(1,33) = 39.55, P < 0.0001; AAV effect: F(1,33) = 21.44, P < 0.0001; *post hoc* test: stress within GFP: ***P < 0.001, within KD: **P < 0.01; AAV effect within CVS: ***P < 0.001. **H**-**N**) CVS-induced dendritic spine changes are reversed by KD of *Ttr*. **H**) Representative images of dendrites. Scale bar 10 µm. **I**) Stubby spines: n = 29-41 from 3-4 mice; AAV effect: F(1,138) = 4.84, P < 0.05; interaction: F(1,138) = 18.64, P < 0.0001; *post hoc* test: stress within GFP: ***P < 0.001, AAV within naïve: ***P < 0.001. **J**) Neck-containing spines: n = 30-40 from 3-4 mice; stress effect: F(1,139) = 14.28, P < 0.001; AAV effect: F(1,139) = 7.86, P < 0.01; interaction: F(1,139) = 92.21, P < 0.0001; *post hoc* test: stress within GFP: ***P < 0.001, within KD: ***P < 0.001; AAV within naïve: ***P < 0.001, within CVS: ***P < 0.001. **K-N**) Cumulative head diameter. **K**) CVS within GFP: χ^2^ = 8.30, df = 1, **P < 0.01. **L**) CVS within KD: χ^2^ = 2.96, df = 1, P = 0.09. **M**) AAV within naïve: χ^2^ = 0.02, df = 1, P = 0.90. **N**) AAV within CVS: χ^2^ = 1.56, df = 1, P = 0.21. **D**-**G**, **I**, **J**) Individual data points are plotted and means **±** s.e.m. are shown. Non-significant comparison not listed. Sketches were generated with biorender.com.

### Altered DNAme on the *Ttr* promoter mediates gene expression changes and behavior

Despite its clinical relevance, *Ttr* gene regulation remains poorly understood. We investigated a 438 bp region proximal to the transcription start site (**Fig. 4A**), focusing on DNA methylation, which is directly regulated by the one-carbon metabolism or indirectly affected via histone methylation.

**Fig. 4:**
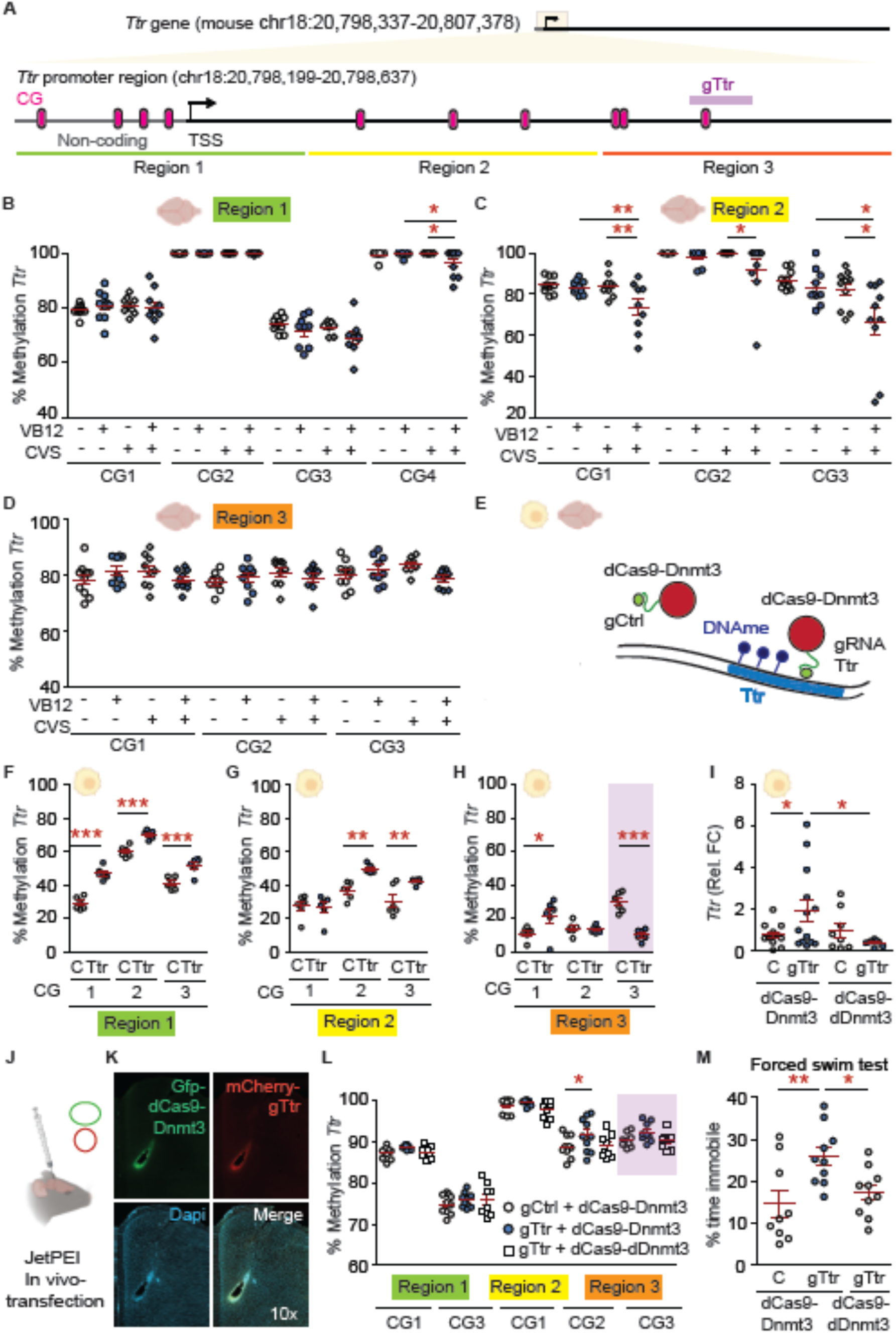
DNAme near the *Ttr* promoter is functionally linked to stress-associated behavior. **A**) Overview of the investigated region. Covered CGs are labelled in pink and refer to listed CGs per region. Regions 1-3 were covered separately by pyrosequencing. **B**-**D**) Pyrosequencing on PFC from mice that had undergone CVS (vs. naïve) and acute vitamin B12 treatment (“VB12”, vs. saline). n = 8-10. **B**) Region 1. CG4: interaction CVS & vit. B12: F(1,32) = 4.17, P < 0.05; drug within CVS: *P < 0.05; stress within VB12: *P < 0.05.. **C**) Region 2. CG1: stress effect: F(1,35) = 7.16, P < 0.05; drug effect: F(1,35) = 5.05, P < 0.05; interaction: F(1,35) = 4.75, P < 0.05; drug within CVS: **P < 0.01; stress within VB12: **P < 0.01; CG2: drug effect: F(1,34) = 4.35, P < 0.05; drug within CVS: *P < 0.05; CG3: stress effect: F(1,36) = 7.37, P < 0.05; drug effect: F(1,36) = 6.08, P < 0.05; drug within CVS: *P < 0.05; stress within VB12: *P < 0.05. **D**) Region 3. CG3: all comparisons n.s. **E**-**M**) Epigenome editing of DNAme on the *Ttr* promoter using a Dnmt3A/L-dCas9 construct (v.s. dDnmt3A/L-dCas9). **E**) Overview. **F**-**I**) Editing in Neuro2A-cells. **F**-**H**) Pyrosequencing. n = 6. **F**) Region 1. gRNA effect: F(1,30) = 110.0, P < 0.001; position effect: F(2,30) = 168.8, P < 0.001; interaction: F(2,30) = 4.01, P < 0.05; *post hoc* test: gRNA effect for all CGs: *** P < 0.001. **G**) Region 2. gRNA effect: F(1,30) = 15.89, P < 0.001; position effect: F(2,30) = 21.7, P < 0.0001; interaction: F(2,30) = 4.96, P < 0.05; *post hoc* test: gRNA effect for CGs 2, 3: ** P < 0.01. **H**) Region 3. Interaction gRNA & CG: F(2,30) = 18.44, P < 0.001; *post hoc* test: gRNA effect for CG1: * P < 0.05, for CG3: *** P < 0.001. **I**) qPCR shows a selective increase in *Ttr* expression after editing. n = 7-13. results. Enzyme effect: F(1,37) = 4.33, P < 0.05; interaction: F(1,37) = 4.33, P < 0.05. *Post hoc* test: Enzyme effect within gTtr: *P < 0.05; gRNA effect within Dnmt3-dCas9: *P < 0.05; **J**-**M**) *In vivo* epigenome editing using jetPEI-transfection. **J**) Overview. **K**) Confocal image of injection site (10x magnification). **L**) Editing alters DNAme across CGs that were not at 100% methylation in the gCtrl-group. n = 8-10; treatment effect: F(2,123) = 6.64, P < 0.01; position effect: F(4,123) = 353.6, P < 0.0001; *post hoc* test: CG2 on region 3: treatment: *P < 0.05. **M**) Increased immobility time in the forced swim test after editing *Ttr*: n = 9-10. One-way ANOVA: F = 5.75, P < 0.01; Tukey’ *post hoc* test: gCtrl guideRNA within Dnmt3-dCas9: **P < 0.01; Enzyme-group within *Ttr*: *P < 0.05. Purple: predicted binding site of gTtr. Individual data points are plotted and means **±** s.e.m. are shown. Non-significant comparison not listed. Sketches were generated with biorender.com.

We utilized pyrosequencing with primers spanning three adjacent regions to analyze *Ttr* DNAme after CVS and vitamin B12 treatment (**Fig. 4B-D**). While the baseline DNAme was high in this region, we detected a vitamin B12-induced reduction in DNAme within the stressed group. To evaluate the functional significance of these observed DNAme changes, we employed epigenome editing with a dCas9-DNMT3ACD-DNMT3LCD-3xFLAG fusion construct (“dCas9-Dnmt3”) along with guide-RNAs for *Ttr* ^14,46^. Twelve gRNAs spanning the three regions were tested in Neuro2A-cells for their impact on *Ttr* gene expression and DNAme (data not shown). A scrambled, non-binding guide RNA (“gCtrl”) was used as a control. The gRNA that most reliably induced DNAme changes while also affecting *Ttr* expression (“gTtr”) was chosen for further analysis (**Fig. 4A**, **Fig. 4E**-**I**). gTtr increased DNAme in multiple assessed CG sites (**Fig. 4F, G**), while reducing DNAme in its binding region (**Fig. 4H**), underscoring the importance of including controls for sterical hindrance during binding. Accordingly, while the combination of gTtr with dCas9-Dnmt3 elevated *Ttr* levels, this effect was not observed with a construct expressing an inactive enzymatic group (“dCas9-dDnmt3”) (**Fig. 4I**). These findings suggest that DNAme near the *Ttr* promoter can impact *Ttr* expression *in vitro*, consistent with vitamin B12-mediated effects in stressed mice, although additional factors likely contribute to mediate stress-induced *Ttr* changes in mice that did not receive vitamin B12 (**Fig. 2**, **Fig. 1B**-**D**).

To evaluate whether DNAme plays a functional role in stress-associated behavioral changes, we combined epigenome editing with *in vivo* jetPEI transfection using constructs labeled with fluorophores for confocal tracking (“gTtr-mCherry” and “dCas9-Dnmt3-GFP”, **Fig. 1J-M**). While we obtained a fluorescent signal in the PFC, only a limited number of cells near the injection site were infected (**Fig. 4K**, compared with viral approaches in **Fig. 2C**, **Fig. 3C**). Consequently, we observed a modest yet significant effect of CRISPR treatment on DNAme across CG sites that were not fully methylated within controls (**Fig. 4L**). The effect was absent with the mute enzymatic construct. Moreover, we noted a significant impact on the immobility time in the forced swim test (**Fig. 4M**), indicating that increased DNAme on the *Ttr* promoter causally contributes to mood- and stress-associated behavior.

## Discussion

In this study, we unveil a tissue- and sex-specific pathway that mediates the acute effects of vitamin B12 supplementation. By employing the chronic variable stress (CVS) mouse model, which exhibits the highest transcriptional-level compliance with depressed patients, we identified *Ttr* as a target affected by vitamin B12. We observed altered *TTR* in the postmortem PFC of male depressed patients. However, it remains unclear whether vitamin B12 supplementation exerts rapid effects on stress response in humans as well and we encourage further investigation in this regard. Lifestyle interventions, particularly those involving dietary factors, are likely to play a more substantial role in personalized medicine. Micronutrients such as vitamins offer well-defined molecular structures with clear biological functions, making them easy to study and administer. In parallel with pre-clinical research, assessment of dietary states and deficiencies in patients may facilitate the treatment of stress-related disorders and guide future clinical studies on the effects of vitamin B12.

The observation that vitamin B12 exerts effects in female mice without altering *Ttr* suggests the involvement of additional mechanisms. In previous studies, we identified other vitamin B12-mediated changes associated with stress, including alterations in *Ntrk-2* and GMPPB signaling. These changes were not detected by the RNA-sequencing performed here, suggesting a regulation on protein levels (e.g. GMPPB methylation) or could be attributed to modified protocols (such as the combination of CMS and acute stress for *Ntrk-2*, which was rescued by vitamin B12). Our cumulative findings point towards a multi-level effect of vitamin B12 and emphasize the need for further research, especially in female cohorts.

While vitamin B12 supplementation is typically associated with an increase in methyl donors, our observation of reduced DNAme within the stress group suggests an indirect effect, possibly through histone methylation. This warrants further investigations in the future. Through using the innovative combination of epigenome editing and jetPEI *in vivo* transfection, we achieved *in vivo* DNAme editing for the first time, enabling direct causal examination of behavioral outcomes. We observed significant alterations in DNAme and behavior.

Through RNA-sequencing following viral overexpression of *Ttr*, we identified a diverse array of transcripts downstream of *Ttr*. Given that *Ttr* is conventionally considered an extracellular protein, the mechanisms through which it regulates these genes remain enigmatic. It is conceivable that the extracellular Ttr protein binds to receptors that initiate transcription cascades. *Ttr* is associated with oxidative stress and can mediate cytotoxicity through Ca^2+^ efflux from the endoplasmic reticulum. Thereby, enhanced Ttr levels could affect neurons within the PFC on multiple levels that may in turn alter gene expression as observed by RNA-sequencing after viral OE of *Ttr*. Alternatively, *Ttr* RNA may interact with intracellular targets. The implicated mechanisms, including immune response, signal transduction, and structural gene products, as well as their impact on depressive-like behavior and neural morphology, should be investigated further.

## Acknowledgments

This study was supported by an Advanced Medical Scientist Award from the Interdisciplinary Center for Clinical Research (IZKF) at the Jena University Hospital, Germany (#973685). Furthermore, this project was made possible by funding from the Carl-Zeiss Foundation (IMPULS #P2019-01-0006). We thank Dr. Thorsten Trimbuch (Charitè Viral Vector Core Berlin, Germany) for input on AAV experiments and Sabine Grunauer-Vasconez (Friedrich-Schiller-University, Jena, Germany) for help with dendritic spine analysis.

## Contributions

G.S., J.S.A., L.L., I.H. and O.E. performed mouse experiments and analysis. G.S., L.L. and O.E. planned experiments and handled paperwork involving animal experiments. G.S. double-checked data and statistics. S.C. and O.E. conducted cell culture experiments. J.S.A., O.E. and A.M. analyzed dendritic spines. E.A.H. provided input on *in vivo* transfection and epigenome editing. K.R. and S.H. analyzed the RNA-sequencing data. The team of T.E. provided help with pyrosequencing. G.T. provided human postmortem samples. C.A.H. provided infrastructure and conceptual advice and edited the manuscript. O.E. designed and planned the study, obtained funding, and wrote the article.

## Supplementary figures

**Supplementary Fig. 1:**
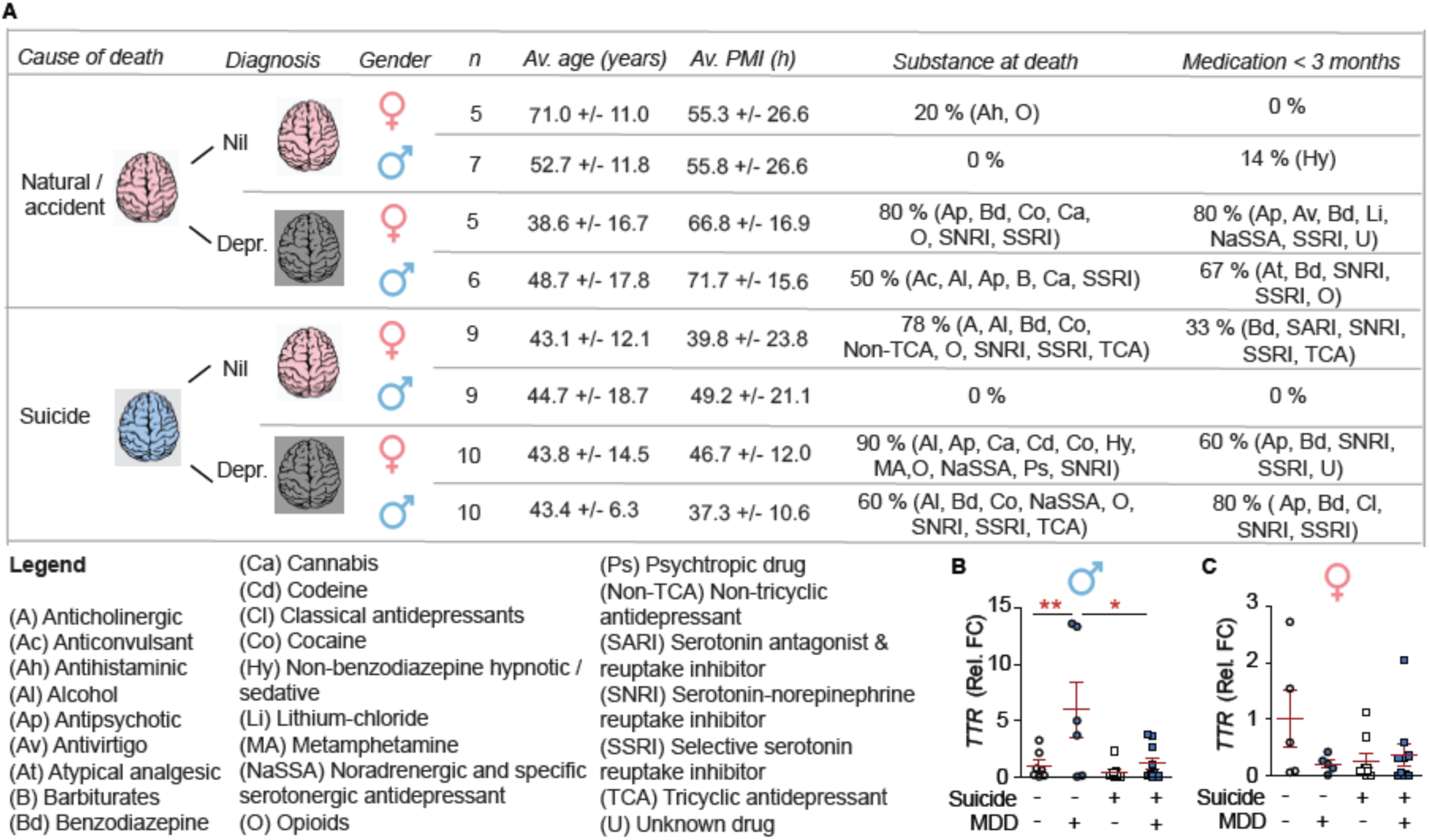
Demographics of postmortem samples and separation by cause of death. **A**) Demographics of tested samples. **B**) Males: n = 7-10; Effect of depr: F(1,28) = 7.94, P < 0.01; effect of cause of death: F(1,28) = 6.85, P < 0.05; interaction: F(1,28) = 4.18, P = 0.05; *post hoc* test: effect of Depr. within non-suicide: **P < 0.01, within suicide: P > 0.05; effect of suicide within healthy (Nil): P > 0.05, within Depr: *P < 0.05. **C**) Females: n = 5-10; effect of depr: F(1,25) = 1.81, P = 0.19; effect of cause of death: F(1,25) = 1.19, P = 0.27; interaction: F(1,25) = 3.26, P = 0.08. **B, C**) Individual data points are plotted and means **±** s.e.m. are shown. Sketches were generated with biorender.com.

**Supplementary Fig. 2:**
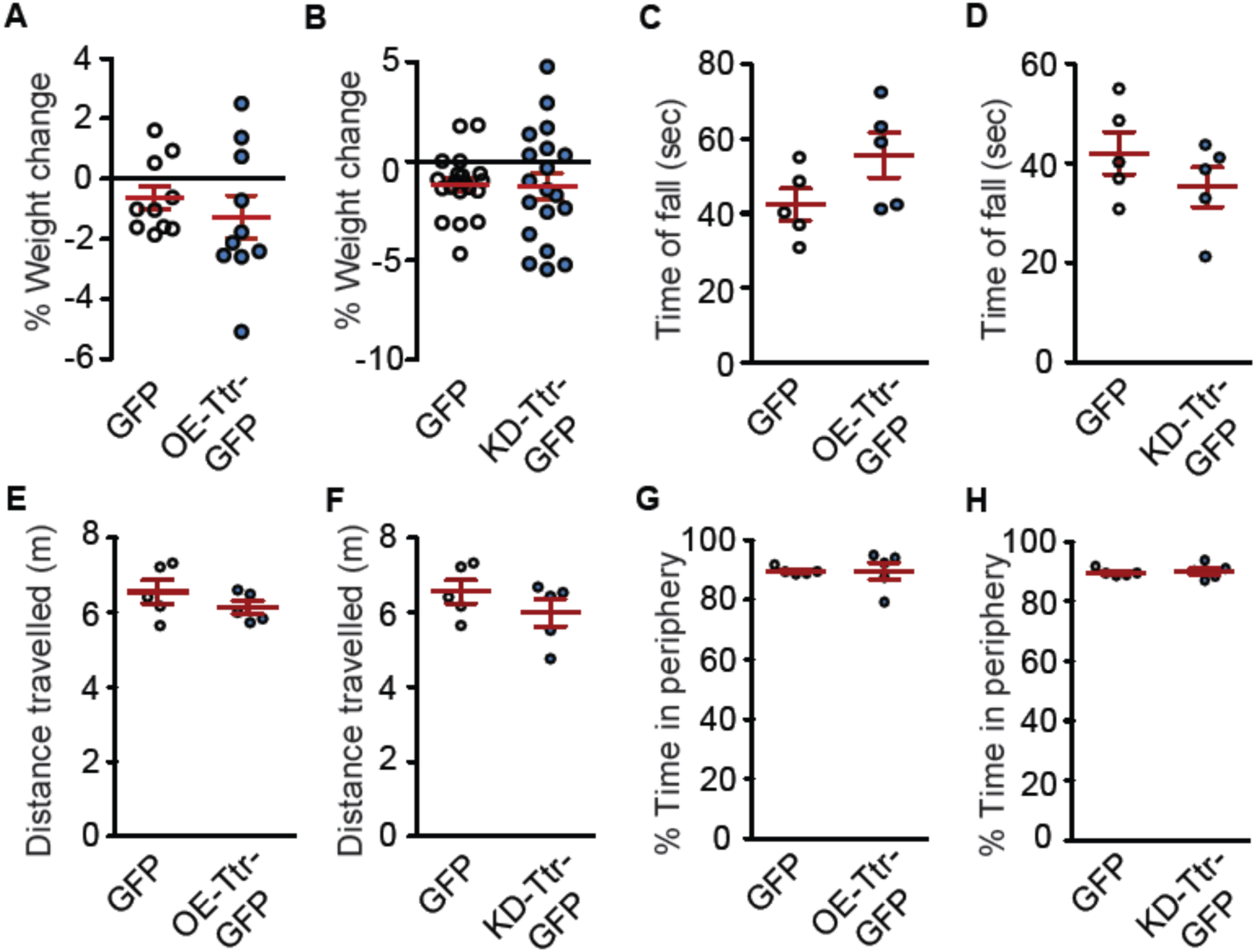
AAVs that alter *Ttr* do not affect body weight or behavior in the open field test. **A**, **B**) % weight change 4 weeks after surgery (before CVS) vs. day of surgery (prior). **A**) Ttr-OE: n = 10; t_18_ = 0.78, P = 0.45. **B**) Ttr-KD: n = 19-20; t_37_ = 0.09, P = 0.93. **C**, **D**) Rotarod, time of fall. **C**) Ttr-OE: n = 5; t_8_ = 1.79, P = 0.11. **D**) Ttr-KD: n = 5; t_8_ = 1.16, P = 0.28. **E**-**H**) Open field. **E**) Ttr-OE: total distance traveled. n = 5; t_8_ = 1.17, P = 0.28. **F**) Ttr-KD: total distance traveled. n = 5; t_8_ = 1.18, P = 0.27. **G**) Ttr-OE: % time in the periphery. n = 5; t_8_ = 0.01, P = 0.99. **H**) Ttr-KD: % time in the periphery. n = 5; t_8_ = 0.34, P = 0.75. **A**-**H**) Individual data points are plotted and means **±** s.e.m. are shown.

**Supplementary Fig. 3:**
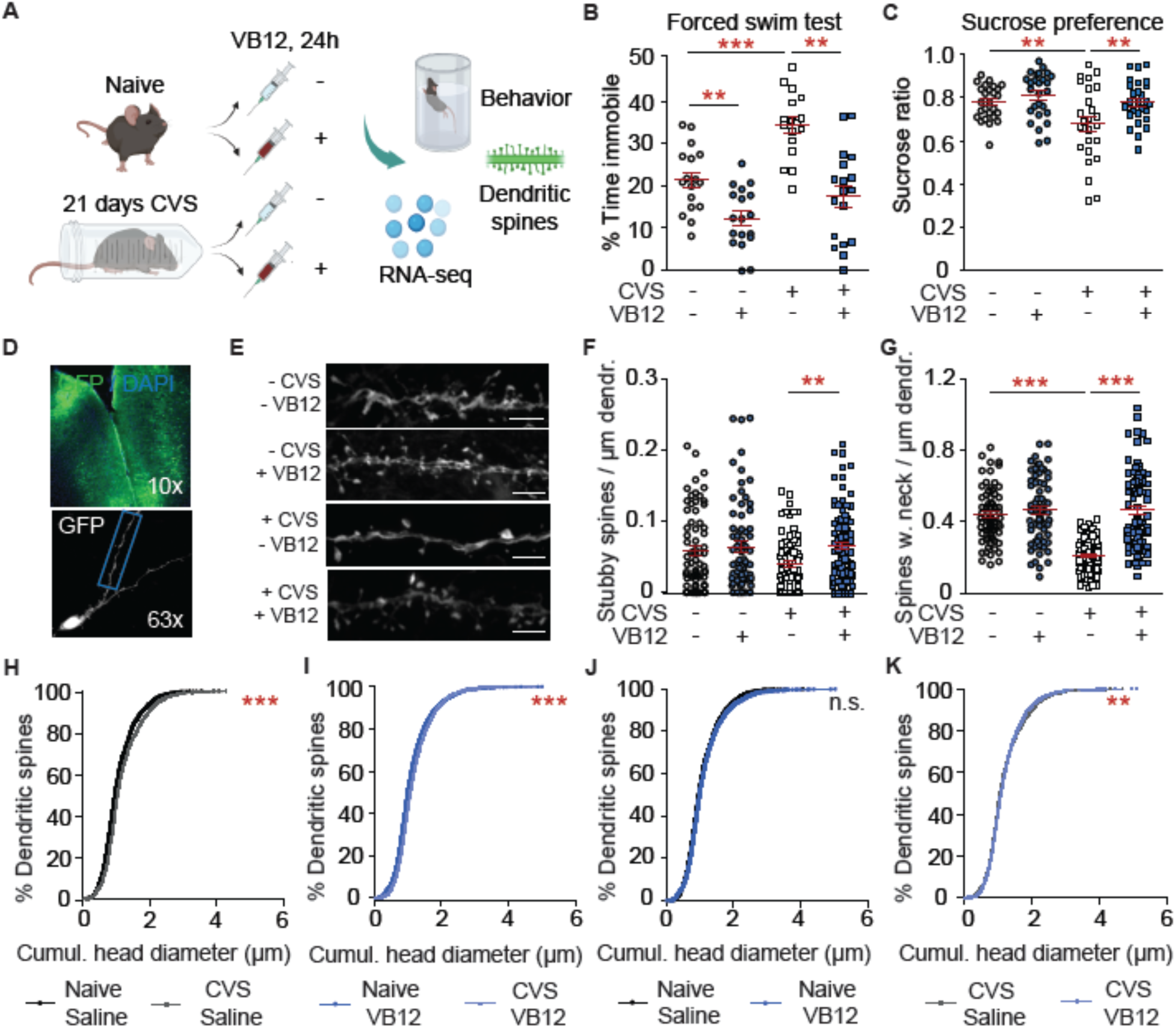
Acute vitamin B12 reverses chronic stress-induced effects on behavior and dendritic spines in the PFC. **A**-**K**) Both sexes were combined. **A**) Experimental plan. **B**, **C**) Stress-induced behaviors are reversed by VB12. **B**) Forced swim test: n = 17-18; stress effect: F(1,66) = 19.95, P < 0.0001; drug effect: F(1,66) = 40.43, P < 0.0001; *post hoc* test: stress within sal: ***P < 0.001; drug within naïve: **P < 0.01, within VB12: ***P < 0.001. **C**) Sucrose preference test: n = 25-28; stress effect: F(1,102) = 7.90, P < 0.01; drug effect: F(1,102) = 7.90, P < 0.01; *post hoc* test: stress within sal: **P < 0.01; drug within VB12: **P < 0.01. **D**-**K**) Dendritic spine analysis. VB12 reverses chronic stress effects. **D**) Overview of AAV-GFP-labelled neurons and dendrites in the PFC. The blue box highlights a secondary dendrite as used for spine analysis. **E**) Representative spines. Scale bar 10 µm. **F**) Stubby spine density: n = 71-89 dendrites from 7-9 mice; drug effect: F(1,302) = 6.88, P < 0.01; *post hoc* test: drug effect within VB12: **P < 0.01. **G**) Density of neck-containing spines: n = 72-89 dendrites from 7-9 mice; stress effect: F(1,306) = 35.14, P < 0.0001; drug effect: F(1,306) = 52.06, P < 0.0001; interaction: F(1,306) = 36.63, P < 0.0001; *post hoc* test: stress within naïve: *** P < 0.001; drug within VB12: ***P < 0.001. **H**-**K**) Cumulative head diameter. **H**) Stress within sal: χ^2^ = 28.47, df = 1, ***P < 0.0001. **I**) Stress within VB12: χ^2^ = 102.7, df = 1, ***P < 0.0001. **J**) Drug within naïve: χ^2^ = 0.53, df = 1, P = 0.47. **K**) Drug within CVS: χ ^2^ = 8.88, df = 1, **P < 0.001. **B**, **C**, **F**, **G**) Individual data points are plotted and means **±** s.e.m. are shown. Non-significant comparisons: not stated. Sketches were generated with biorender.com.

**Supplementary Fig. 4:**
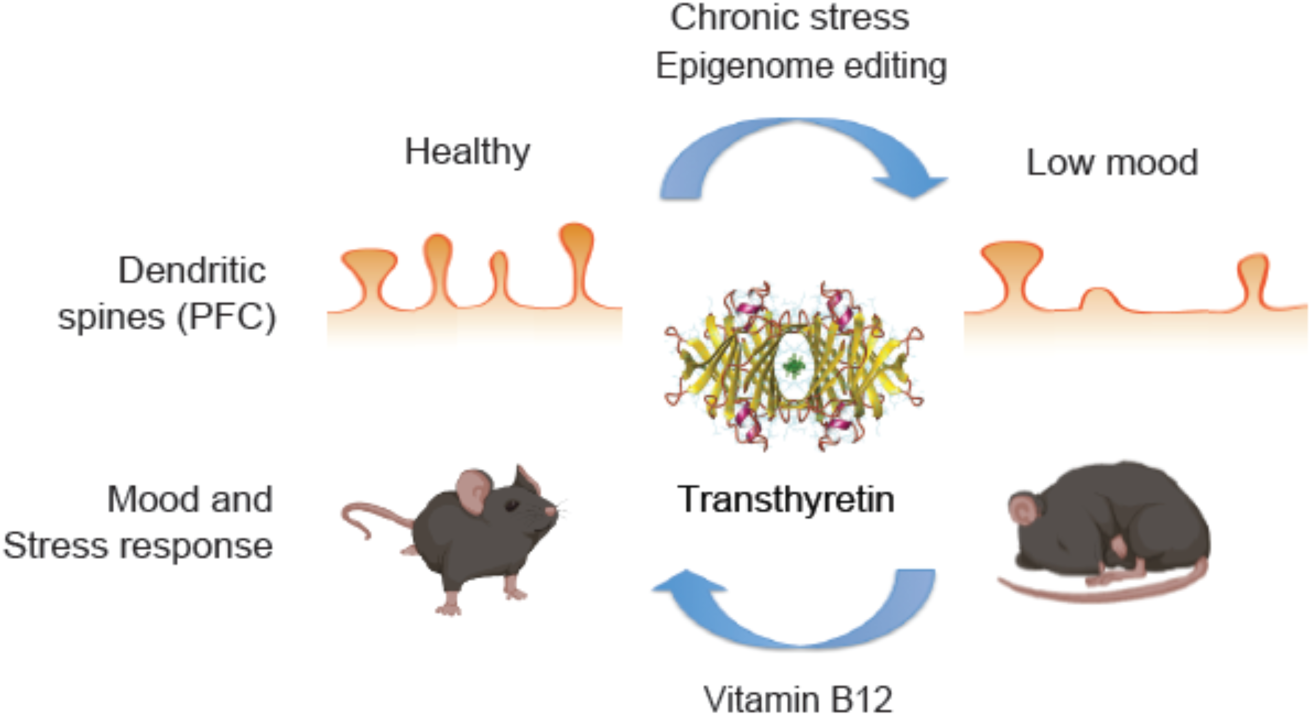
Overview of proposed mechanism.

## Supplementary methods

### 1. Primers

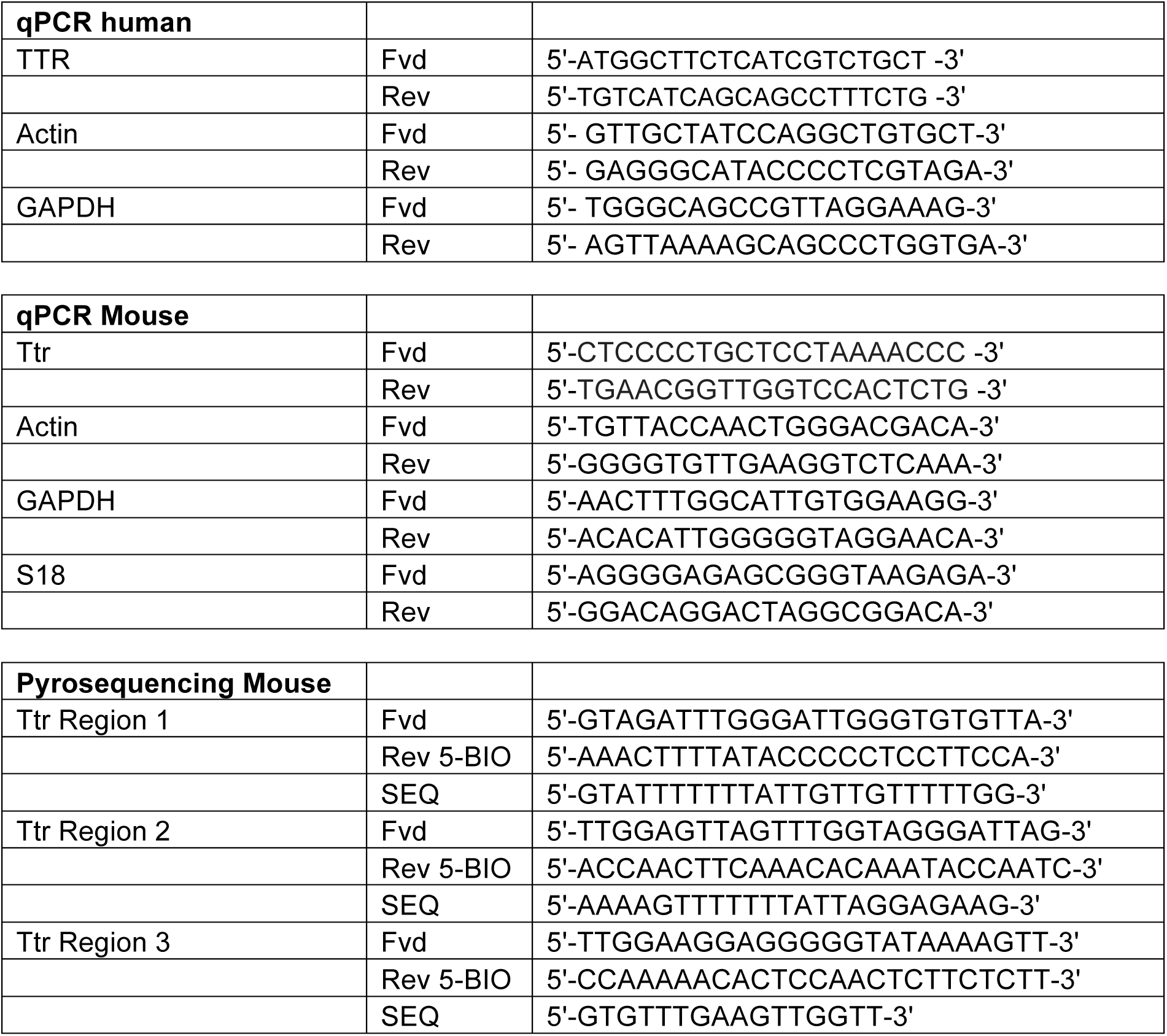

### 2. Extended methods

#### AAVs & stereotaxic surgery

Bilateral stereotaxic surgery into the dorsomedial PFC was essentially performed as described ^17^. Mice received were analgized and anesthesized using 100 mg/kg ketamin (Ketabel 100mg/ml, Bela-Pharm) and 16mg/kg xylazin (Rompun 2%, Bayer Animal Health GmbH). Eyes were protected with Vitamycin cream (#CP3920, CP Pharma). The skin on the skull was opened and small holes were drilled near the stereotaxic X/Z coordinates of the PFC (coordinates relative to Bregma: anteroposterior (Z): + 1.8, mediolateral (X): ±0.8, dorsoventral (Y): -2.75, angle: 15°). Viruses were injected at a rate of .1 µL/min for 5 min using 10 µL Gastight Hamilton syringes model 1801 RN. Mice were scored for general health and behavior at least 3 days post-surgery. The following three viruses were utilized: pAAV.1-CAG-GFP (#37825, Addgene), OE-Ttr-GFP: pAAV-CAG-GFP-P2A-Ttr-WPRE.bgH (custom-made by Charitè viral vector core, Berlin, Germany), KD-Ttr-GFP: pAAV-U6-Ttr-shRNA-2-CAG-GFP-WPRE3 (custom-made by Charitè viral vector core, Berlin, Germany).

#### RNA purification and quantification

RNA was purified by resuspension in 1ml Trizol (Thermo Fisher Scientific, #15596026). After 5 min incubation at room temperature (RT), 200 µl chloroform were added to precipitate the RNA. Samples were shaken vigorously for 15 sec and incubated for 3 min at RT. They were then centrifuged for 15 min at 13.000 rpm at 4°C. The clear part of the upper phase was transferred into a new eppendorf tube. After adding 500 µl isopropanol and mixing by inversion, samples were incubated for 10 min at RT and then centrifuged for 10 min at 13.000 rpm and 4°C. The supernatant was discarded and the pellet was washed with 1ml 75% ethanol. The samples were centrifuged for 5 min at 8.500 rpm at 4°C. For RNA-sequencing, the washing step with ethanol was repeated. The resulting pellet was resuspended in 30-100 µl of ddH_2_O depending on the size of the pellet. Quality and quantity of RNA were assessed by Nanodrop and Qubit. RNA was converted to cDNA using cDNA conversion reagents (#A5001, Promega, Madison, WI, USA) according to the manufactorers’ instructions. Quantitative realtime-PCR was done on a Bio-rad CFX96 Real-time system using SsoFast EvaGreen Supermix (##1725201, Bio-Rad).

#### DNA methylation analysis

For DNA-purification, cell were lysed for at least 30 min at 56°C in lysis buffer (50 mM Tris-HCl pH 8.0, 100 mM EDTA, pH 8.0, 100 mM NaCl, 1% SDS, 0.3mg/ml proteinase K) until all tissue was dissolved. Then, 500 µl of phenol/chloroform/isoamylalcohol (#A156.1, Roth), were added and samples were rotated for 30 min at RT. After centrifugation for 15 min at 14.000 rpm at RT, the entire upper phase was collected into a new eppendorf tube. DNA was precipitated with 50 µl of 8M lithium chloride and 500 µl isopropanol. After mixing by inversion, samples were spun for 15 min at 14.000 rpm at RT and the supernatant was discarded. The pellet was washed with 1 ml of 70% ethanol and spun for 10 min at 14.000 rpm at 4°C. The pellet was resuspended in 30 – 100 µl ddH_2_O, depending on the size of the pellet. Quality and quantity of DNA were assessed by Nanodrop. Bisulfite conversion was performed using the EZ DNA Methylation-Gold kit (#D5006, Zymo, Irvine, CA, USA). PCR was performed using the PyroMark PCR Kit (#978703, Qiagen) according to the manufactorers’ instructions. Annealing temperatures for *Ttr* pyrosequencing primers during PCR were 58°C (region 1 and 3) and 60°C (region 2). Sequencing was conducted on a Pyromark Q96 Sequencer.

